# Glycogen Engineering Improves the Starvation Resistance of Mesenchymal stem cells and their Therapeutic Efficacy in Pulmonary Fibrosis

**DOI:** 10.1101/2025.02.21.639504

**Authors:** Yongyue Xu, Mamatali Rahman, Zhaoyan Wang, Bo Zhang, Hanqi Xie, Lei Wang, Haowei Xu, Xiaodan Sun, Shan Cheng, Qiong Wu

**Affiliations:** State Key Laboratory of Green Biomanufacturing, PR China; MOE Key Laboratory of Bioinformatics, Center for Synthetic and Systems Biology, Tsinghua University, Beijing, 100084, China; School of Life Sciences, Tsinghua University, Beijing, 100084, China; School of Materials Science and Engineering, Tsinghua University, Beijing, 100084, China; Department of Medical Genetics and Developmental Biology, School of Basic Medical Sciences, Capital Medical University, Beijing 100069, China; Xinjiang Stem Cells Special Plateau Disease Engineering Technology Research Center, The First People’s Hospital of Kashi, 66 Yingbin Road, Kashi, 844000, PR China

**Author notes:** Corresponding author. State Key Laboratory of Green Biomanufacturing, MOE Key Laboratory of Bioinformatics, Center for Synthetic and Systems Biology, School of Life Sciences, Tsinghua University, Beijing, 100084, China. These authors contributed equally to this work (Yongyue Xu, Mamatali Rahman).

**Keywords:** Mesenchymal stem cells, Glycogen, Pulmonary Fibrosis, Glycogen synthase, Cell survival

## Abstract

Mesenchymal stem cells (MSCs) are widely used in regenerative medicine, including the treatment of pulmonary fibrosis. However, implanted MSCs disappear within days, constraining therapeutic efficacy, which is largely attributed to nutrient deprivation. In this study, we established glycogen metabolism engineering strategies in mammalian cells. By expressing a functionally optimized glycogen synthase (GYSmut), MSCs could accumulate large amounts of glycogen rapidly as a reserve substance. Glycogen engineering significantly improved the survival of MSCs during starvation both in vitro and in vivo, enhancing cell viability post-implantation and their therapeutic efficacy in pulmonary fibrosis. Glycogen-engineered MSCs may serve as chassis cells for further applications. Our research highlights the importance of glucose metabolism regulation in cell-based therapy and demonstrates the great potential for the metabolic engineering of MSCs and other therapeutic cells.

## Introduction

Pulmonary fibrosis (PF) is traditionally regarded as an irreversible lung disease for which there are generally no effective drugs, with the exception of nintedanib and pirfenidone ^1,2^. Regenerative medicine represents an alternative approach, with several successful therapeutic applications in patients with chronic lung diseases, including PF ^3–5^. The recent focus on mesenchymal stem cells (MSCs) has generated enthusiasm due to their unique regenerative and immunomodulating properties, including the ability to reverse fibrosis ^6,7^. Recent studies have begun to unravel the mechanisms underlying the therapeutic potential of MSCs, with a particular emphasis on their secretory properties. The paracrine factors secreted by MSCs, including fibroblast growth factor and cytokines, have been shown to mediate tissue repair and regeneration ^6, 8^. However, the therapeutic efficacy of MSCs has been limited due to their poor survival and short dwell time in the hostile pathological microenvironment, which has hindered the translation of these promising preclinical findings into clinical practice^9, 10^.

Metabolic regulation is crucial for cell survival post-implantation. It has been reported that the hypoxic in vivo microenvironment inhibits oxidative phosphorylation (OXPHOS) in implanted MSCs, forcing them to rely on glycolysis instead, which yields less ATP ^11–13^. Our previous single-cell transcriptomic study also showed that the glucose metabolism pathway of MSCs was downregulated in the PF microenvironment ^6^. Compared with in vitro culture systems, it is more difficult for implanted MSCs to obtain sufficient glucose in pathological microenvironments, and recent findings suggest that glucose exhaustion is the main reason for cell death after implantation ^14^. Supplying glucose to MSCs improved their survival post-implantation ^15^. Therefore, glucose metabolism engineering has the potential to improve MSC therapy by enhancing starvation resistance and prolonging the residence time of implanted cells.

Glycogen is a branched polymer that acts as the primary glucose storage form in mammalian cells, supplying energy under glucose deficiency and promoting cell survival. Glycogen metabolism is controlled by a series of enzymes, including glycogen synthase (GYS) ^16^. To reduce cell death of implanted MSCs caused by nutrient deprivation, we engineered the glycogen metabolism of MSCs with genes encoding essential enzymes, which promoted glycogen accumulation (Fig. 1A). The accumulated glycogen serves as an energy supply post-implantation, improving the viability and therapeutic efficacy of MSCs (Fig. 1B).

**Fig. 1.**
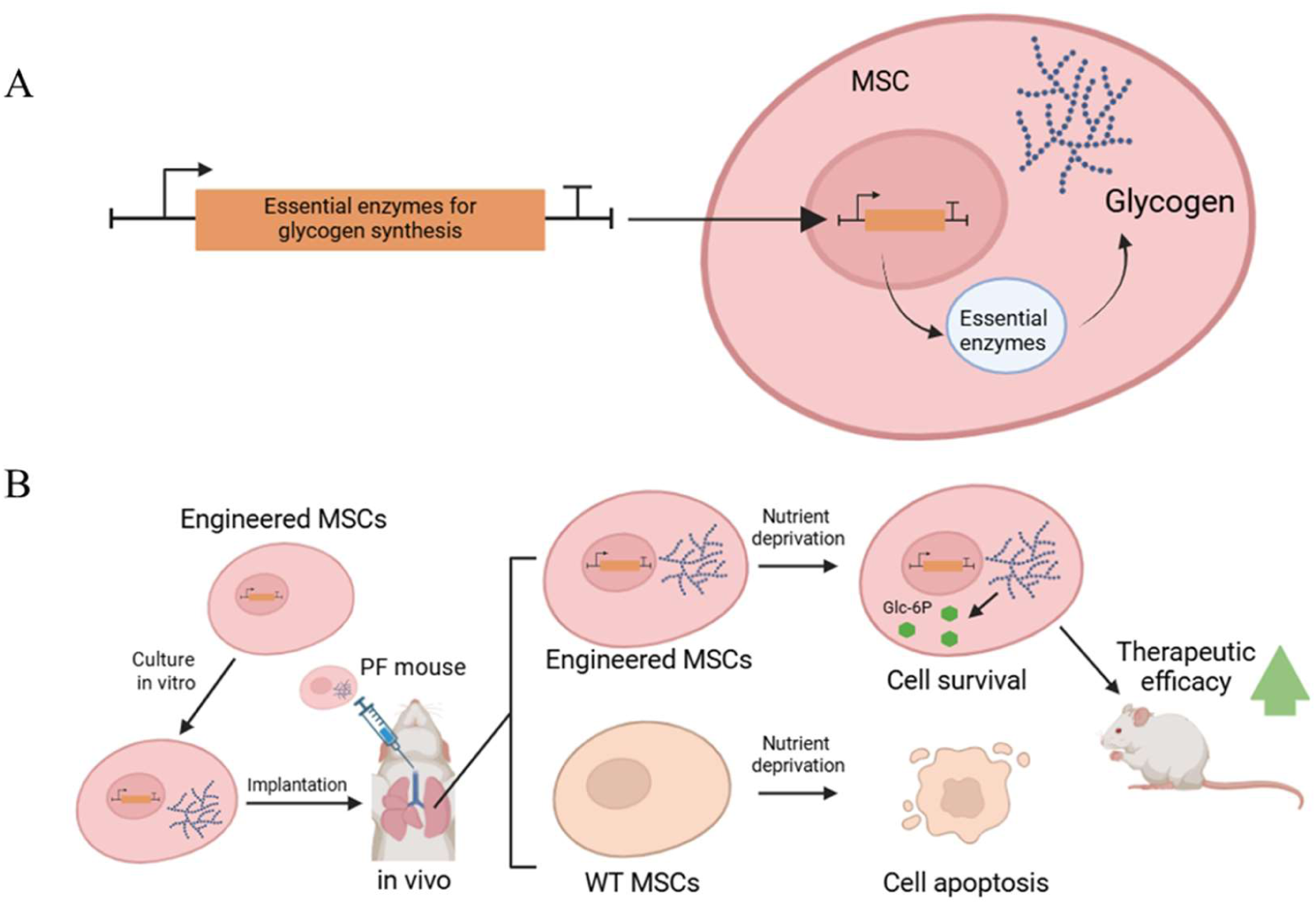
Schematic illustration of the strategy used to enhance the therapeutic efficacy of MSCs through glycogen engineering. (A) Engineering glycogen metabolism of MSCs by overexpressing essential enzymes of glycogen synthesis. (B) Engineered MSCs use glycogen as an energy supply after implantation, improving cell viability and therapeutic efficacy.

## Results

### Construction of glycogen strategies in mammalian cells

Glycogen is synthesized from UDP-glucose in a process involving glycogenin (GYG), UDP-glucose pyrophosphorylase (UGP), glycogen synthase (GYS), and glycogen-branching enzyme (GBE) ^16^. GYG is the initiator of glycogen synthesis, UGP catalyzes the synthesis of UDP-glucose, GYS is responsible for the synthesis of glycogen, and GBE introduces branches into the structure of glycogen (Fig. 2A). Here, we firstly devised glycogen engineering strategies based on these essential enzymes.

**Fig. 2.**
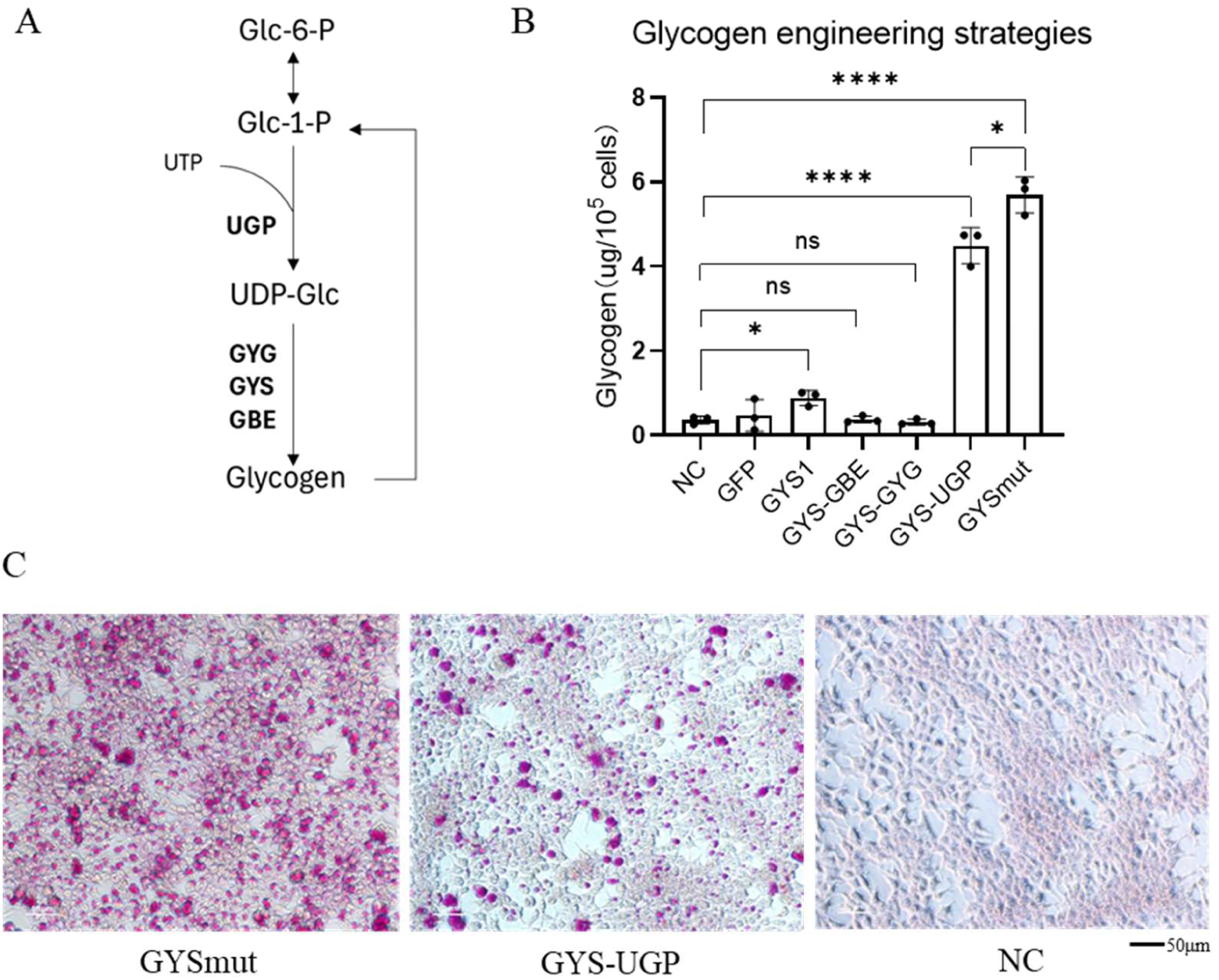
Construction of glycogen engineering strategies. (A) Essential enzymes of glycogen synthesis. (B) Tests of essential enzyme combination strategies in HEK293T cells by transient transfection. The glycogen content of each group was measured. (C) Periodic acid-Schiff (PAS) staining of cells expressing GYSmut and GYS-UGP revealed significant accumulation of glycogen granules.

Initially, we transiently expressed *GYS1* in human HEK293T cells, but it only had a limited, albeit significant effect on glycogen accumulation (Fig. 2B). We attributed the unsatisfactory results to substrate limitation, as well as potential inhibition of *GYS1* by cellular pathways. Next, we tried different expression strategies to increase glycogen accumulation, including (1) expressing *GYS1* alone to promote glycogen synthesis, (2) co-expressing *GYS1* with *GBE1* to increase glycogen-branching, (3) co-expressing *GYS1* with *GYG1* to increase glycogen initiation, and (4) co-expressing *GYS1* with *UGP1* to increase the substrate supply. Some protein kinases (e.g., AMPK) inactivate GYS1 by phosphorylation at specific serine sites ^17^. Therefore, (5) we introduced Ser-Ala mutations at the Ser-8, Ser-641, Ser-645, Ser-649, Ser-653, Ser-654, and Ser-657 sites of GYS1 (GYSmut) to prevent phosphorylation. Both co-expression of *GYS1* with *UGP1* and expression of GYSmut alone resulted in greatly increased glycogen accumulation compared to the control (∼12 and 16 folds) and other strategies (Fig. 2B). Glycogen granules were clearly visible following periodic acid-Schiff (PAS) staining (Fig. 2C).

Since most of the Ser-Ala mutation sites were located near the C-terminus of GYSmut, we truncated the 100 C-terminal amino acids to generate GYSdelc. Although this truncated variant was still capable of inducing cellular glycogen accumulation, it exhibited a reduced rate in comparison to full-length GYSmut (Fig. S1A). Further, we co-expressed GYSmut with UGP, which demonstrated a marked additive effect (Fig. S1B).

### Glycogen accumulation improved the starvation resistance of MSCs

After expressing GYSmut in MSCs by lentivirus transduction, glycogen accumulation was significantly increased in the engineered MSCs (Fig. 3A). PAS staining showed more PAS-positive glycogen granules in GYSmut MSCs (Fig. 3B), which altered cell morphology by significantly increasing the cell volume, as assessed by flow cytometry (Fig. S2A). Glycogen granules were located both in the cytoplasm and nucleus (Fig. S2B). Interestingly, when co-overexpressing of wild-type GYS1 and GYG, we observed a diffuse intracellular glycogen distribution rather than bulky glycogen particles, indicating that the combination of distinct enzymes influences the characteristics of glycogen synthesis (Fig. S2C).

**Fig. 3.**
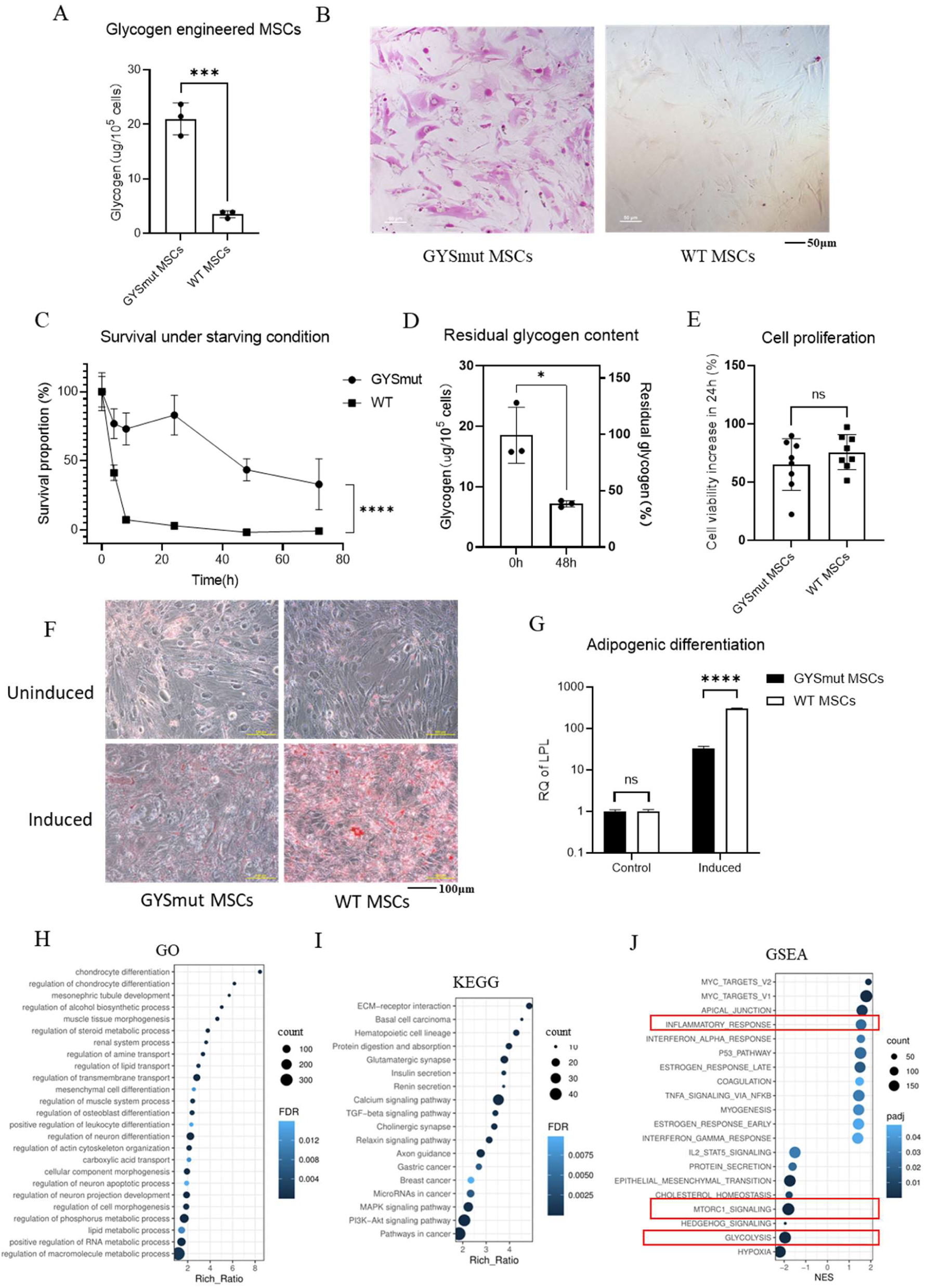
Construction of glycogen-engineered MSCs. (A) Glycogen content of GYSmut MSCs. (B) PAS staining of GYSmut MSCs, showing glycogen granules (red). (C) Survival of engineered MSCs under DPBS (starvation) treatment in vitro (N = 8). (D) Residual glycogen content of GYSmut MSCs after DPBS (starvation) treatment for 48 hours. (E) Viability of GYSmut MSCs according to the CCK8 assay. (F, G) Adipogenic differentiation potential of GYSmut MSCs, assessed by oil red O staining and qPCR detection of Lpl expression (n=3, unpaired *t*-test P-value < 0.0001). (H) GO enrichment analysis of differentially expressed genes (DEGs) between GYSmut MSCs and the GFP control. (I) KEGG analysis of DEGs. (J) Gene set enrichment analysis (GSEA) of the DEGs.

To evaluate the starvation resistance of glycogen-engineered MSCs, we cultured GYSmut MSCs in Dulbecco’s Phosphate-Buffered Saline (DPBS) instead of a complete culture medium to create a nutrient-deprived environment. The Cell-Counting-Kit-8 assay was used to measure cell viability at various time points. Notably, approximately 40% of the GYSmut MSCs were still viable after 72 hours of starvation culture, whereas wild-type MSCs died rapidly after 8 hours (Fig. 3C). After 48 hours of starvation, GYSmut MSCs still retained a 39% glycogen reserve (Fig. 3D), indicating that the accumulated glycogen could be a long-term energy resource to maintain higher cell viability during nutrient deprivation following implantation. Since some pathological contexts are hypoxic, we tested the starvation resistance of MSCs under hypoxia (oxygen concentration < 5%), with the engineered MSCs still maintaining a higher survival proportion (Fig. S2D).

As glycogen is a crucial node of carbon metabolism in mammalian cells, we next investigated whether glycogen engineering caused global changes in MSCs. Cell counting showed that glycogen engineering had no significant effect on cell proliferation (Fig. 3E). To evaluate the effects of glycogen engineering on the adipogenic differentiation capacity of MSCs, we induced GYSmut MSCs with adipocyte medium for 5 days, followed by oil red O staining and qRT-PCR of the adipocyte differentiation marker *Lpl*. The results indicated a decrease in the adipocyte differentiation potential of GYSmut MSCs (Fig. 3F, 3G). It was reported that adipogenic differentiation and the immunomodulatory functions of MSCs are mutually exclusive ^18^, so blocking adipocyte differentiation may enhance the immunosuppressive activity of MSCs.

We next performed bulk RNA sequencing (RNA-seq) to understand the effect of glycogen engineering on the transcriptome of GYSmut MSCs. A total of 759 genes were up- and 542 were downregulated in GYSmut MSCs, indicating a broad effect on the transcriptome (Fig. S2E). Gene ontology (GO) enrichment analysis revealed that differentially transcribed genes were predominantly enriched in processes related to cellular differentiation, morphogenesis, and the regulation of various metabolic pathways, including lipid metabolism, reflecting potential metabolic reprogramming. The differentially expressed genes were also enriched in the terms related to processes of muscle, kidneys, and neural tissue, which are sites where glycogen is naturally distributed in mammals (Fig. 3H) ^16^. KEGG analysis showed the PI3K-Akt signaling pathway was also modulated, which is crucial for cell survival (Fig. 3I). The GSEA results showed that the inflammatory response gene set was upregulated, which may be beneficial in the treatment of PF, while mTORC1 and glycolysis-related gene sets were downregulated (Fig. 3J). It was reported that mTORC1 responds to cellular amino acid and glucose levels ^19^, which is in agreement with the hypothesis that glycogen synthesis affects cellular glucose levels and the process of glycolysis.

### Glycogen engineering promoted the survival of MSCs post-implantation

In our previous study, intratracheal administration of MSCs exhibited a robust therapeutic effect in a mouse model of bleomycin-induced pulmonary fibrosis (PF) ^6^, and further scRNA-seq analysis of implanted MSCs within the PF microenvironment revealed downregulation of glucose metabolism (Fig. S3), suggesting that nutrient availability is a critical determinant of MSC viability and may potentially limit their therapeutic efficacy. We, therefore, tested whether glycogen engineering would promote the survival of GYSmut MSCs in vivo. PF mice were intratracheally injected with glycogen-engineered MSCs co-expressing GYSmut and *Gaussia* luciferase (GYSmut-Gluc), while the control group was injected with Gluc MSCs. We collected lungs of PF mice on days 0 and 7 after MSC implantation, and measured the fluorescence intensity of lung tissue homogenates with a commercial kit, to assess the counts of living MSCs (3 mice each group each time point) (Fig. 4A). A higher proportion of surviving GYSmut-Gluc MSCs was detected than in the control (Fig. 4B).

**Fig. 4.**
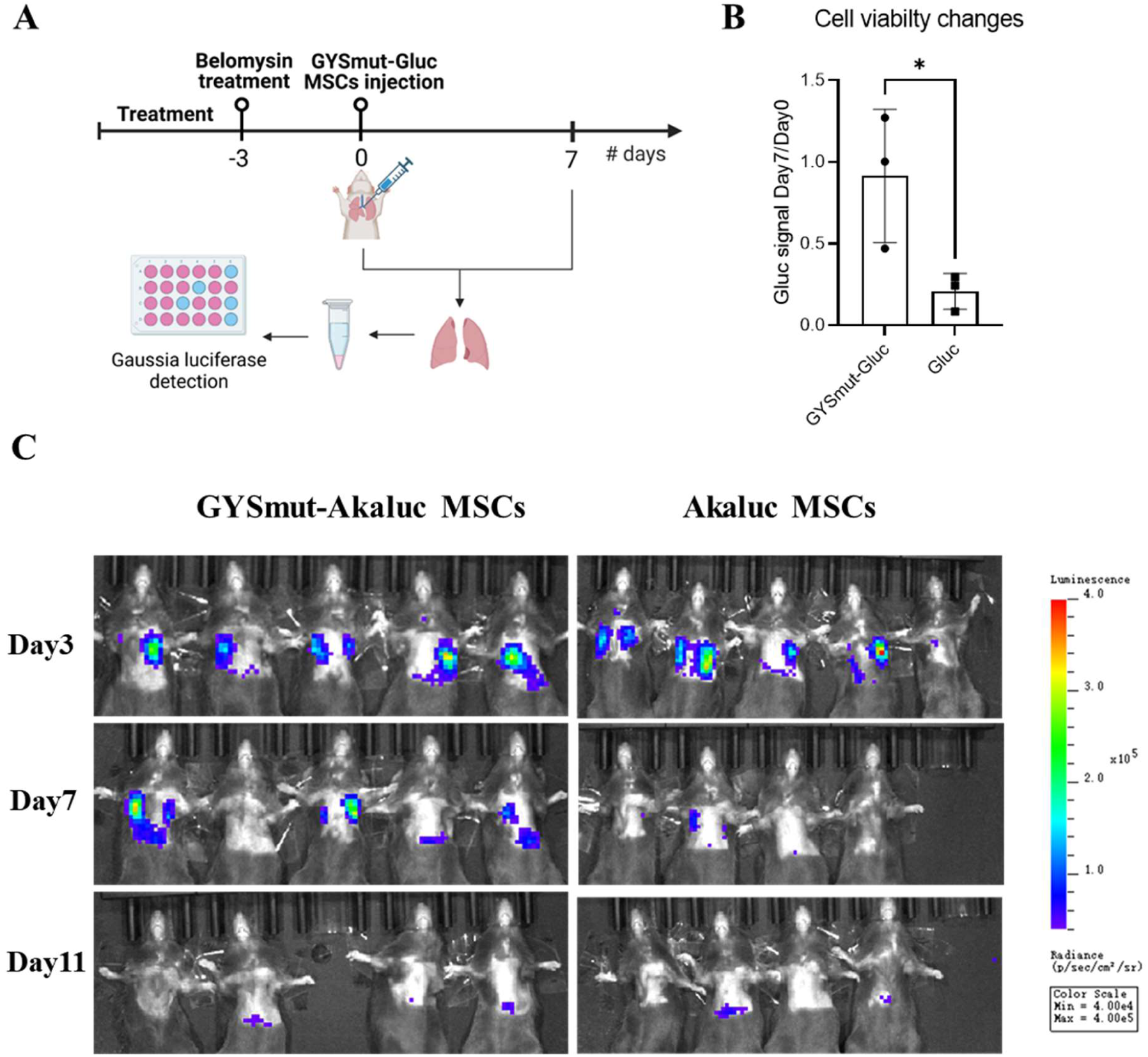
Survival of implanted glycogen-engineered MSCs. (A) Schematic illustration of the strategy used to assess the survival of MSCs by detecting *Gaussia* luciferase activity in the homogenate of lungs. (B) Changes of *Gaussia* luciferase activity in the two groups on day 7 post-implantation (3 mice each group, unpaired *t*-test P-value = 0.044). (C) Live imaging of *Akaluc* luciferase activity in the two groups implanted with GYSmut-Akaluc MSCs and control cells (5 mice each group). One mouse from the control group died on day 7, and one mouse from the GYSmut-Akaluc group died on day 11.

Because of the limited sensitivity of *Gaussia* luciferase labeling for *in vivo* imaging, to assess the persistence and biodistribution of implanted MSCs within the PF microenvironment, we used a different luciferase, *Akaluc* luciferase. GYSmut MSCs co- expressing Akaluc (GYSmut-Akaluc) were intratracheally injected into PF mice. This approach, in combination with the substrate Akalumine ^20^, allows for highly sensitive live imaging and thus provides insights into the retention of implanted cells. In the control group, MSCs expressing Akaluc were injected (5 mice each group). As expected, on day 7 post-implantation, higher fluorescence intensity was detected in the GYSmut MSCs group compared with the control (Fig. 4C, Fig. S4), indicating the greater survival ability of GYSmut MSCs compared to wild-type MSCs in the PF microenvironment. Quantification of fluorescent imaging revealed a further decline in survival rates to similar levels in both GYSmut MSCs and the control by day 11, suggesting that glycogen engineering appears to be primarily functional during the first two weeks post-implantation (Fig. S4).

### Engineered MSCs demonstrated a beneficial effect on pulmonary fibrosis

To investigate the therapeutic efficacy of glycogen-engineered MSCs (GYSmut MSCs), we administered them via tracheal injection into mice on day 3 following bleomycin treatment, and recorded the survival rate and body weight of mice over time (10 mice each group). GYSmut MSCs demonstrated a more pronounced effect than the control group expressing only GFP (Fig. 5A). Quantitative analysis of lung tissue sections stained with H&E and Masson’s trichrome revealed that the GYSmut MSCs group exhibited superior therapeutic efficacy on day 7, as evidenced by reduced collagen deposition and preserved alveolar size (quantified by mean linear intercept, MLI) compared to the control (Figs. 5B and C). The improved therapeutic efficacy illustrated the beneficial effect of enhanced MSC viability and confirmed the rationale for glycogen engineering.

**Fig. 5.**
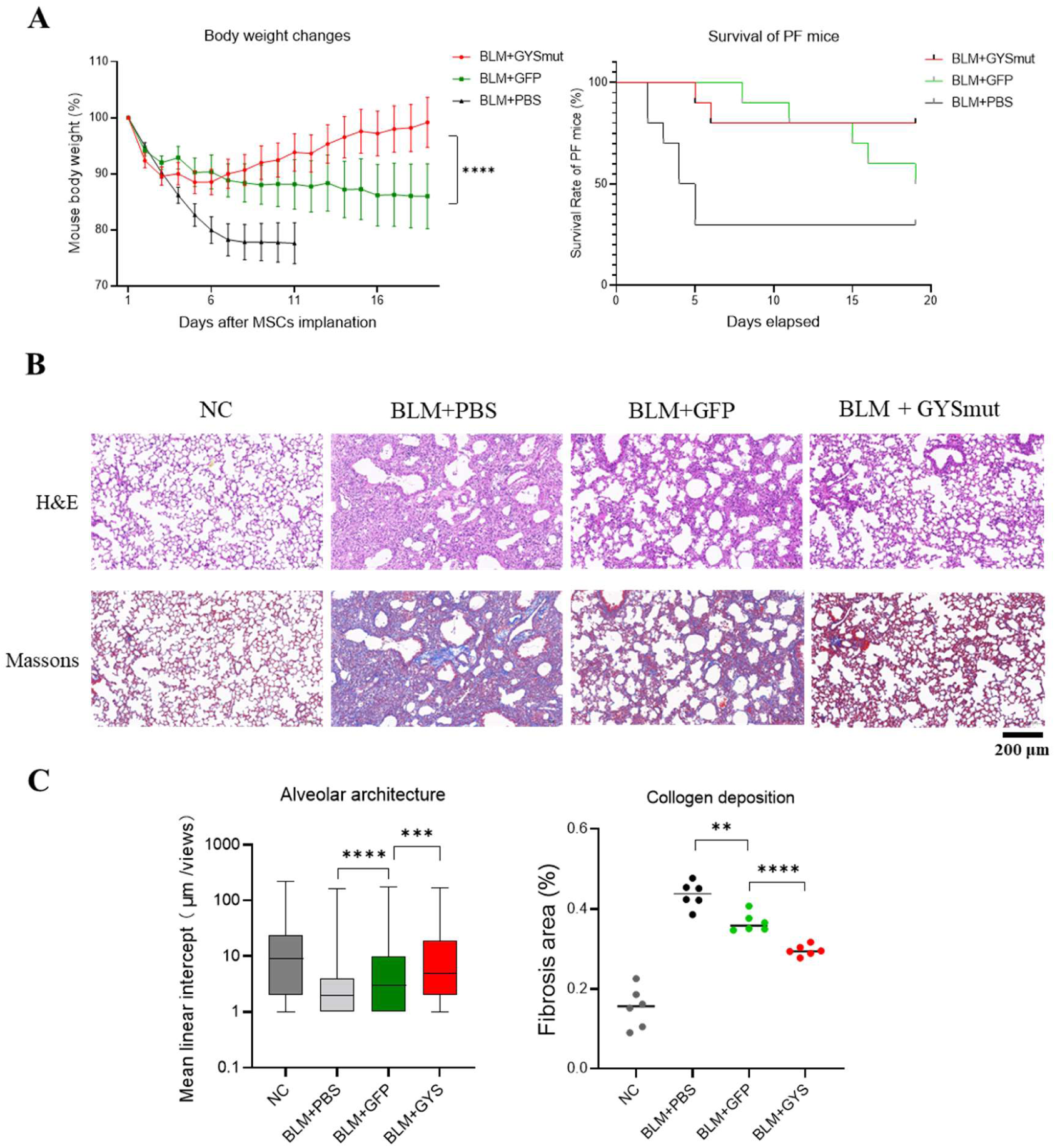
Therapeutic efficacy of glycogen-engineered MSCs. **(A)** Survival and body weight changes of PF mice treated with GYSmut MSCs and control cells (10 mice each group, mean ± SEM, 2-way ANOVA P-value=0.0001), (**B**) Representative lung tissue sections stained with H&E and Masson’s trichrome. NC group is healthy mice. **(C)** Collagen deposition and preserved alveolar size (quantified by mean linear intercept, MLI) of lung tissue sections (6 mice each group).

## Discussion

Glycogen is found in a wide range of organisms, ranging from bacteria to mammals. As a major form of stored glucose, glycogen plays an essential role in maintaining energy homeostasis and cell survival. In this study, we engineered the glycogen metabolism of MSCs to promote glycogen accumulation. We tested several engineering strategies in human HEK 293T cells and primary MSCs derived from mice. Over-expression of phosphorylation-resistant GYS (GYSmut), as well as the co-expression of GYS and UGP, induced intense glycogen. Glycogen engineering enhanced the starvation resistance of MSCs and improved their viability post-implantation. Compared with the control group, the engineered MSCs exhibited a significantly enhanced therapeutic effect on pulmonary fibrosis, indicating the importance of glucose metabolism regulation for MSC-based therapy.

Glycogen-engineered MSCs may serve as chassis cells for further applications by enhancing cell viability post-implantation, and our glycogen engineering strategies also have the potential to be applied to other therapeutic cells. Transient expression of GYSmut by mRNA transfection may be a safer alternative than lentiviral transduction, avoiding the risks of chromosomal modification. LNP-mediated circular mRNA transfection would further improve engineering efficiency^21^.

We investigated the effect of glycogen engineering on the transcriptome of MSCs by RNA-seq, which revealed large-scale gene transcription changes. In future studies, complex regulatory networks may be elucidated at the levels of proteomics and metabolomics. Previous studies have revealed roles of glucose metabolism regulation in MSCs’ immunoregulatory properties ^22, 23^. The broader impact of glycogen engineering on MSCs remains to be elucidated. We tested the therapeutic efficacy of glycogen-engineered MSCs in the bleomycin-induced PF mice model. Their efficacy in other disease models is still to be investigated.

Within the context of MSC-based cellular therapies, administered cells were distributed to various microenvironments characterized by variable oxygen concentration, nutrient availability, and immune responses. While *in vivo* animal models offer a more physiologically relevant platform for validation, they are less convenient for detailed investigation due to multifactorial influences, making it challenging to fully understand the precise kinetic properties and adaptation dynamics of the implanted cells, which is a limitation of this study.

In previous studies, complex gene circuits were introduced to enhance the efficacy of various cell-based therapies, including MSCs and CAR-T cells ^24, 25^, and some research also revealed the advantages of metabolic engineering (ME) ^26^. However, in contrast to extensive ME research in prokaryotic organisms, ME of mammalian therapeutic cells has yet to be developed. Here, we demonstrated the great potential of glycogen engineering, offering a basis for future studies. By dynamically controlling essential enzymes, cellular metabolism can be remodeled to achieve optimal survival and therapeutic efficacy post-implantation. Further research in this area will provide additional opportunities for cell therapy, gene therapy, and tissue engineering.

## Materials and methods

### Animals and experimental design

All the animal experiments were authorized by the institutional animal care and use committee of Tsinghua University. Six-week-old male C57BL/6J mice were housed in the specific pathogen-free experimental animal environment at the laboratory animal research center of Tsinghua University, with 5∼6 mice per cage having access to sterilized food and water ad libitum under a 12-hour light/dark cycle. Mice were anesthetized with avertin (200μL per mouse) or 2.5% tribromoethanol. And euthanized by cervical dislocation or CO2 inhalation. 76 mice were used in total. To obtain reliable results, each group contains at least 3 mice. In the same experiment, the mice in each group came from the same batch and were randomly captured, processed and assigned to different cages. The number of mice is randomly assigned, the surgeon knows the grouping situation, and the statistical analyst does not know the grouping in advance. All animal experiments were authorized by the institutional animal care and use committee of Tsinghua University (Approval No. 23-WQ1.G23-1, Study on different factors on the survivability of implanted MSCs in murine pulmonary fibrosis model, 2023-01-18 to 2026-04-12).

### Isolation of mouse adipose-derived stem cells

MSCs were obtained by euthanizing the mice, digesting the inguinal adipose tissue with collagenase type I (1.0 mg/ml, Sigma), and culturing the released cells in mouse ADSC basal medium (CYAGEN, China) in a humidified atmosphere containing 5% carbon dioxide at 37°C, according to a previously published method ^6^.

### Construction of the BLM-induced PF mouse model and administration of MSCs

Mice were anesthetized with 2.5% tribromoethanol (0.8% NaCl, 1 mM Tris (pH 7.4), 0.25 mM EDTA (pH 7.4)) and administered 3.5 mg/kg bleomycin solution (Sigma, B5507-15un) in 50 μL phosphate-buffered saline (PBS) intratracheally on day 0 using a 1 mL syringe with a 25G needle. Control animals were mock-treated with PBS ^6^.

Injection of MSCs (at passages 3∼6) was performed on day 3 after BLM treatment. Each animal received an intratracheal injection of 50 μL of a suspension comprising 1 × 10^7^ cells/mL in PBS (5 × 10^5^ cells/mouse) using a 1 mL disposable syringe with a 25G needle. For the control and BLM + PBS groups, 50 μL PBS was injected instead. Before implantation, the cells were washed three times to remove the culture medium. The body weight of treated mice was recorded daily ^6, 27^.

### Histological evaluation of lung damage and collagen deposition of BLM-induced lung injury

To assess the development of PF and therapeutic efficacy, mice were evaluated on day 7. For lung tissue collection, mice were euthanized by CO2 inhalation. The thoracic cavity was opened and the lungs were perfused and washed with ice-cold PBS. The lungs were then fixed with 4% (v/v) paraformaldehyde in PBS and processed into 5-μm-thick paraffin sections, which were mounted on glass slides.

For histological assessment of alveolar architecture and fibrosis, sections were stained with hematoxylin and eosin (H&E) or Masson’s trichrome stain, respectively. The samples were scanned using Pannoramic SCAN (3DHISTECH) with a 20× objective. Fibrosis area percentages and alveolar size (quantified by mean linear intercept, MLI) were calculated according to a previously published method^6, 28^.

### Generation of genetically engineered MSCs and in vitro tests

HEK293T cells were cultured in DMEM with 10% FBS. Sequences of essential enzymes are amplified from cDNA of mouse MSCs. To test combinations of essential enzymes, HEK293T cells in 6-well plates were transiently transfected with encoding plasmids (2μg/well) using Lipo8000 (Beyotime, C0533). Co-expressed enzymes were connected by the 2A sequence. After 3 days, the glycogen content of each group was quantified using a commercial kit (SOLARBIO, BC0345-100T). A PAS staining kit (Beyotime, C0142S) was used to stain glycogen granules.

To construct the lentiviral vectors, designed gene sequences were cloned into the pCDH backbone and used to co-transfect HEK293T cells with pMD2.G and psPAX2 plasmids using Lipo8000. Essential enzymes were co-expressed by connecting them with EGFP or puromycin N-acetyltransferase via the 2A sequence. The cell supernatant was collected every 24 hours for 3 days. The viral particles were concentrated using Lentivirus Concentration Solution (Servicebio, G1801-100ML), after which the infectious solution was prepared by resuspending them in 200μL PBS and stored at -80℃ until use. When the density of MSCs in the 6-well plate reached approximately 80%, 20 µl of the lentiviral solution was added. Following 2 days after infection, puromycin was added to a final concentration of 1 mg/L for selection.

Glycogen was quantified after 3 days of induction. To simulate starvation conditions, DPBS solution was used to replace the culture medium of seeded MSCs (96-well plate, 5000 cells/well), and a CCK-8 assay kit (BIORIGIN, BN15201) was used to measure cell viability. To test the adipogenic differentiation potential of MSCs, we induced them with commercial adipocyte medium (Procell, PD-027) for 5 days as described in the manufacturer’s protocol, and then performed oil red O staining (Beyotime, C0157S).

### Viability detection and live imaging of MSCs post-implantation in the PF model

To detect viability changes of MSCs post-implantation, 2.5 × 10^5^ MSCs expressing *Gaussia* luciferase suspended in 50 µl of PBS were injected intratracheally into PF mice, as described above. The whole lungs of mice were collected on the day of surgery (D0) and the 7^th^ day (D7) after surgery (n=3 each time point), and were stored in a 1.5 mL centrifuge tube at -80 ℃. Then, 400 μL PBS and 20 1-mm-beads (Servicebio, G0201) were added to the centrifuge tube with lungs for processing with a KZ-III high-speed homogenizer (Servicebio) at 60Hz for 30s, 6 times. The homogenate was centrifuged (10000g, 5 min) and the activity of *Gaussia* luciferase in the suspension was detected using a commercial kit (GeneCopoeia, LF061). The lungs of cells without Gaussia luciferase expression were used as negative controls, and the corresponding signals were considered to represent background subtraction in the calculations. Cell viability changes of each group were assessed by calculating the ratio of the intensity of lung tissue collected on D7 to D0.

To detect the in vivo distribution of MSCs post-implantation, MSCs expressing Akaluc were injected intratracheally into PF mice (n=5), as described above. Live imaging was performed on the 3^rd^ and 7^th^ days after implantation. Akalumine-HCL (InvivoChem, V41343) was dissolved in PBS (2.5 mg/mL) and intraperitoneally injected into mice (100 μL/mouse). Luminescence was detected after 10 minutes using a Lumina III instrument and analyzed with Living Image software version 4.8.

### RNA-seq of MSCs

MSCs were cultured in 100-mm dishes (5×10^6^ cells per dish) in DMEM/F-12 medium. Trizol (Vazyme, R401-01) was used for RNA extraction. Illumina RNA-seq and subsequent analysis were performed by ANNOROAD Corporation (Beijing, China). DESeq2 was used to identify differentially expressed genes (|log2 fold-change|≥1, adjusted p-value < 0.05).

### Statistical analysis

Statistical analysis was performed using GraphPad Prism software version XY (GraphPad Software Inc.). Significant differences between groups were identified using unpaired Student’s *t*-test or 2-way ANOVA. Asterisks were used to indicate significance in figures (*: p<0.05, **: P<0.01, ***: p<0.001, ****: p<0.0001). Unless indicated otherwise, all quantitative data are presented as mean values ± SD.

## Authors’ contributions

Y.Y.X. and M.R. designed the experiments. Q.W. supervised the project. Y.Y. X., M. R., Z.Y. W., H.W. X. and B.Z. performed the experiments. H.Q. X., L. W., and S. C. analyzed the data. Q.W., Y.Y.X., M. R. and X.D. S. wrote the manuscript.

## Acknowledgments

We thank the team of Du Yanan at Tsinghua University for sharing the plasmid encoding Akaluc. We also thank Prof. George Guo-Qiang Chen at Tsinghua University for his advice on this project.

The authors declare that they have not used Artificial Intelligence in this study.

## Statements and Declarations Ethical considerations

All animal experiments received approval from the institutional animal care and use committee of Tsinghua University (Project title: Study on different factors on the survivability of implanted MSCs in murine pulmonary fibrosis model, Approval number: 23-WQ1.G23-1, Date of approval: 2023-01-18).

## Availability of data and materials

The dataset generated by this study is available in GSE281974 [https://www.ncbi.nlm.nih.gov/geo/query/acc.cgi?acc=GSE281974]. Additional data that support the findings of this study are available from the corresponding author upon reasonable request.

---

**Fig. S1.**
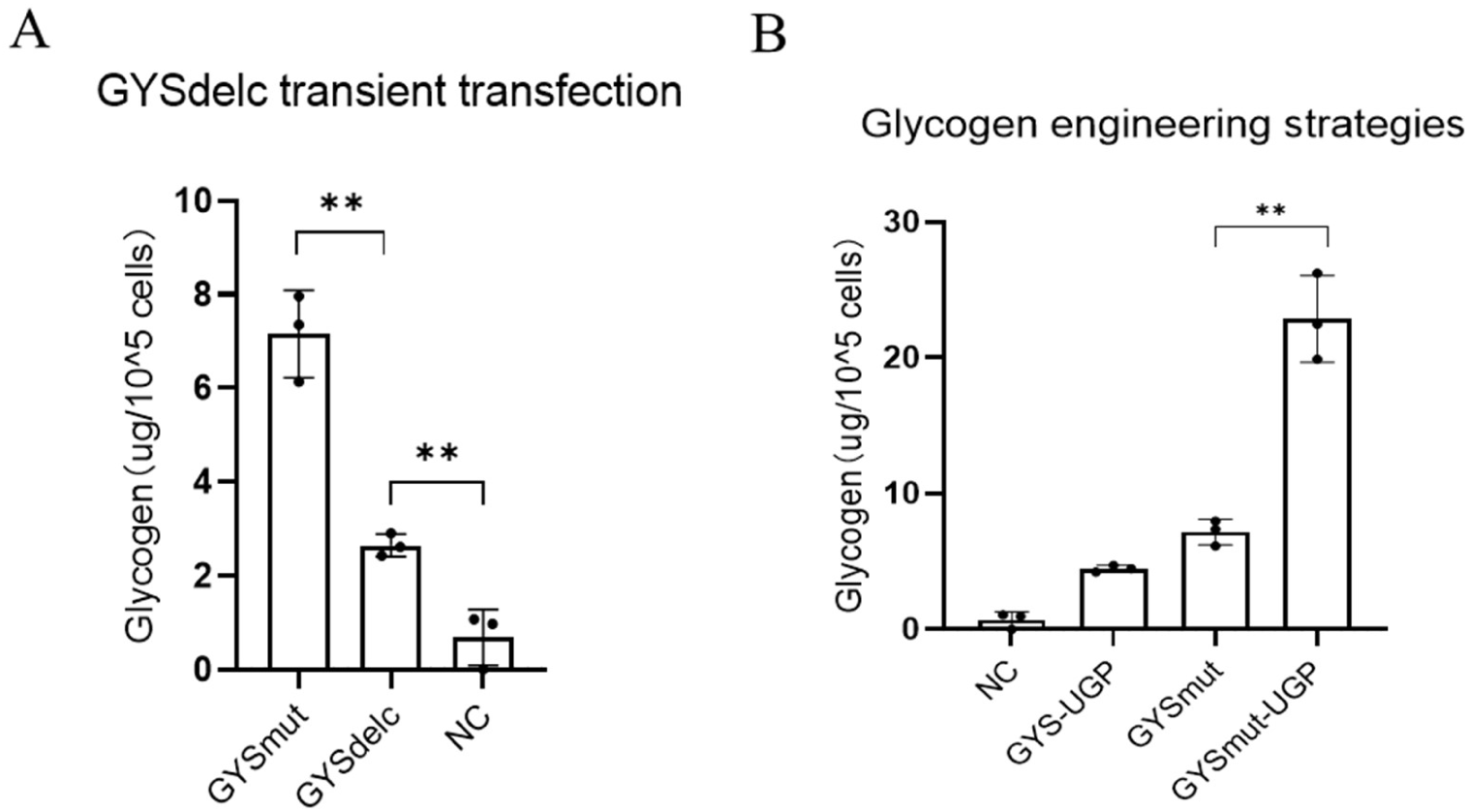
(A) GYSdelc promoted glycogen accumulation. (B) Co-expressing GYSmut and UGP further promoted glycogen accumulation.

**Fig. S2.**
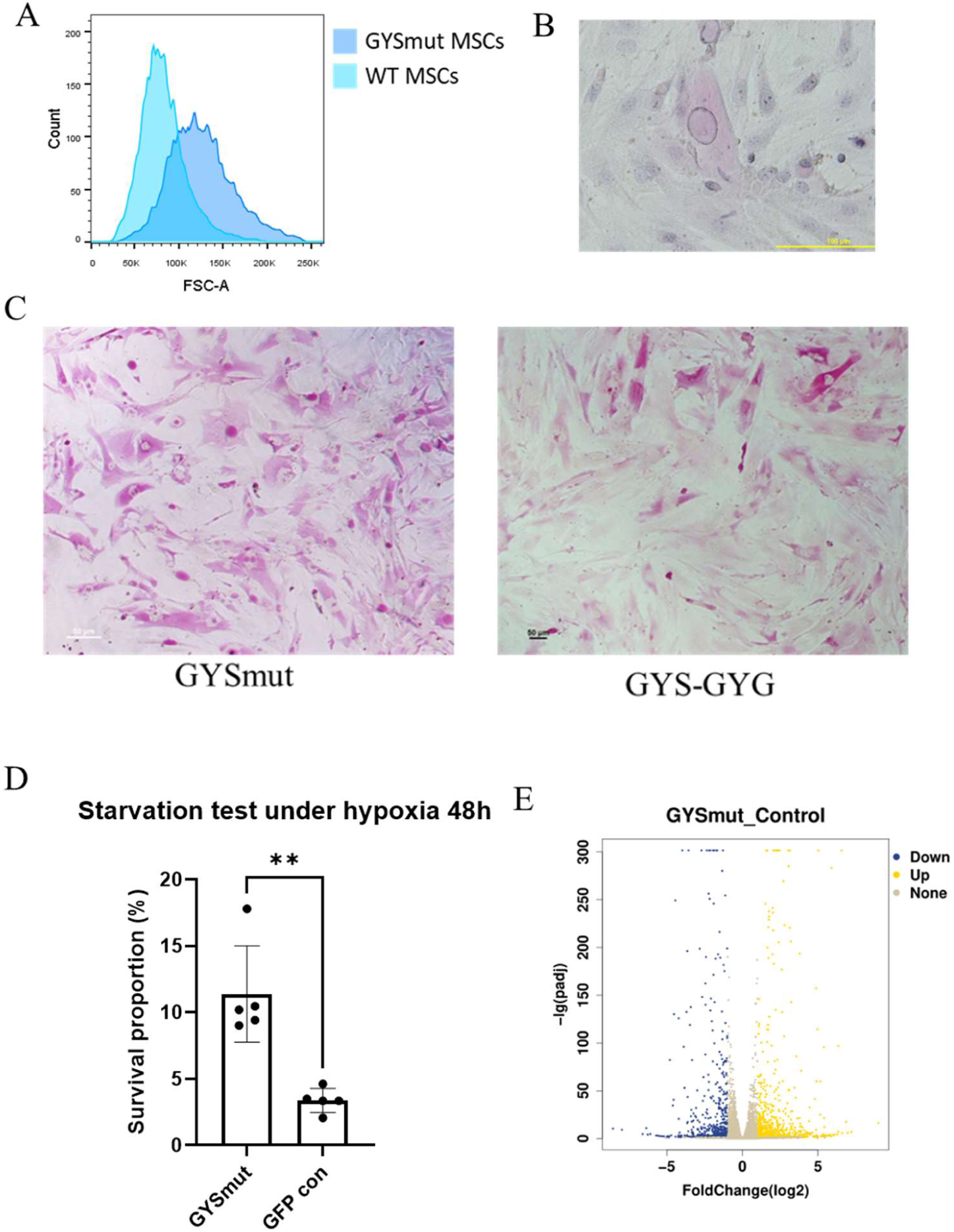
Impacts of glycogen engineering on MSCs. (A)Cell volume change of engineered MSCs, assessed through flow cytometry. (B) PAS and hematoxylin staining of GYSmut MSCs. Glycogen (red) is distributed in cell nucleus (blue) and cytoplasm. (C)Glycogen distribution of GYSmut and GYS-GYG MSCs. Scale bar, 50 μm. (D)Starvation resistance test under hypoxia. (E) Impacts of glycogen engineering on transcriptome of MSCs.

**Fig. S3.**
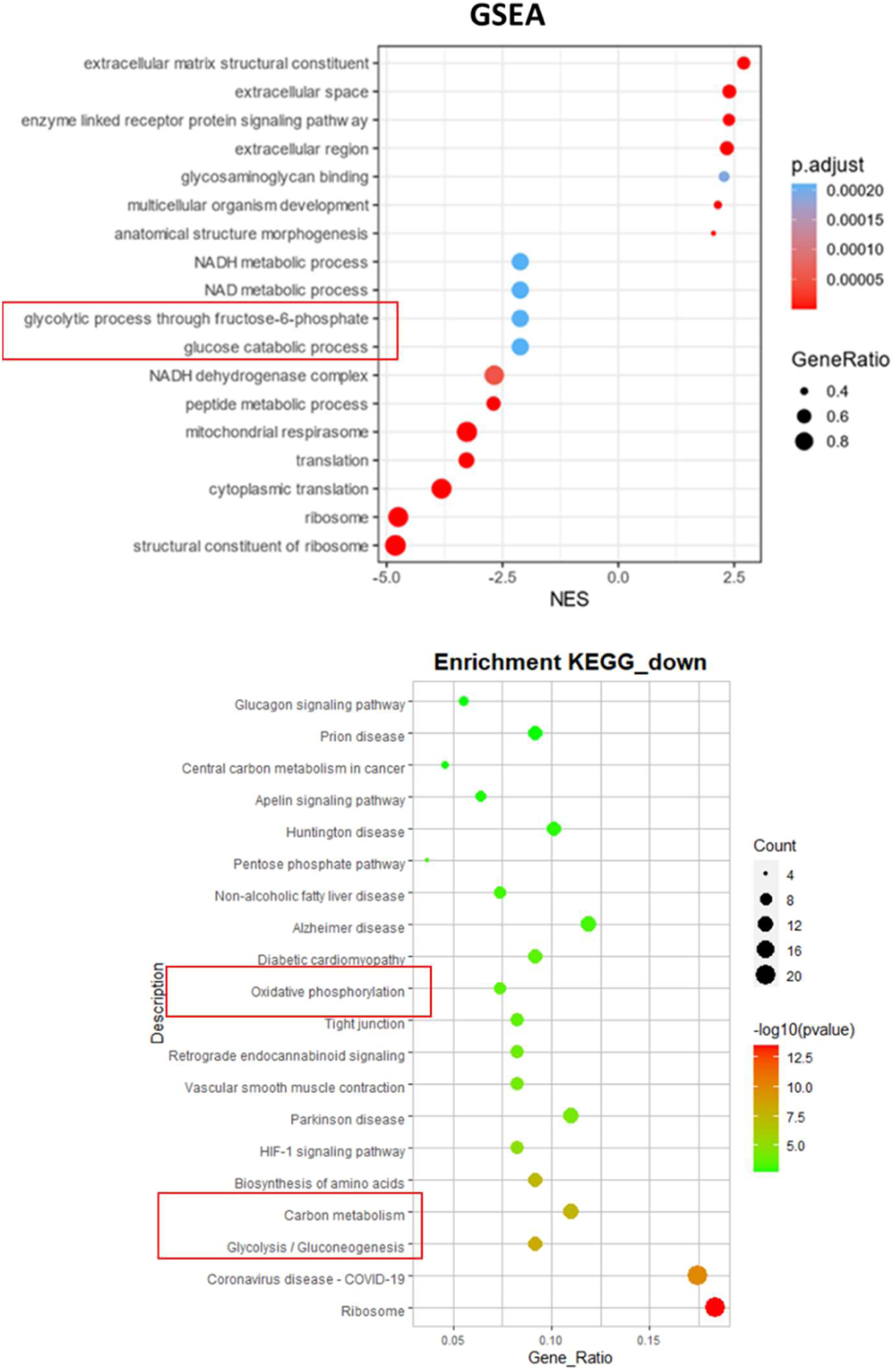
KEGG and GSEA analysis of implanted MSCs in our previous research. GFP MSCs were intratracheally administered to PF mice and collected through flow cytometry. Single-cell sequencing showed downregulation of glucose metabolism.

**Fig S4.**
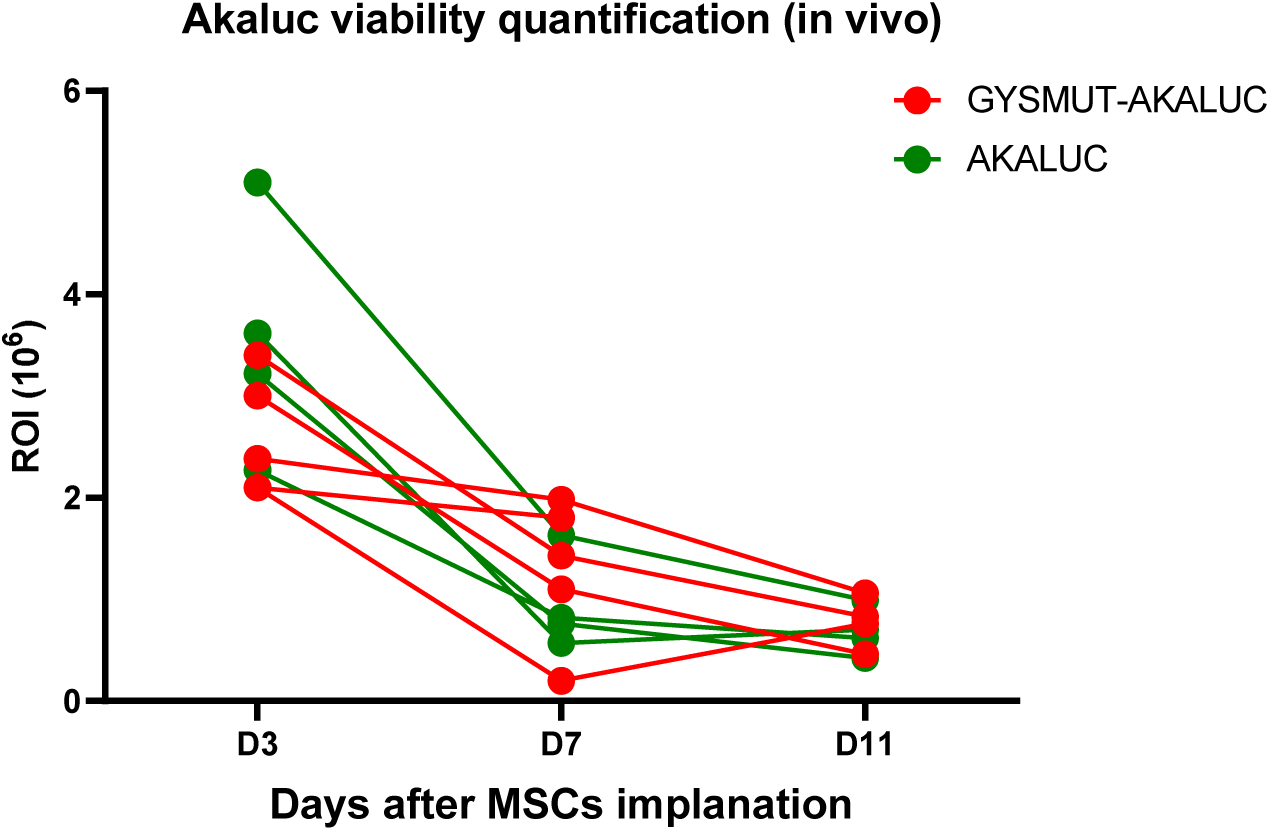
Quantification of akaluc activity of in vivo imaging post-implantation.

**Table 1:**
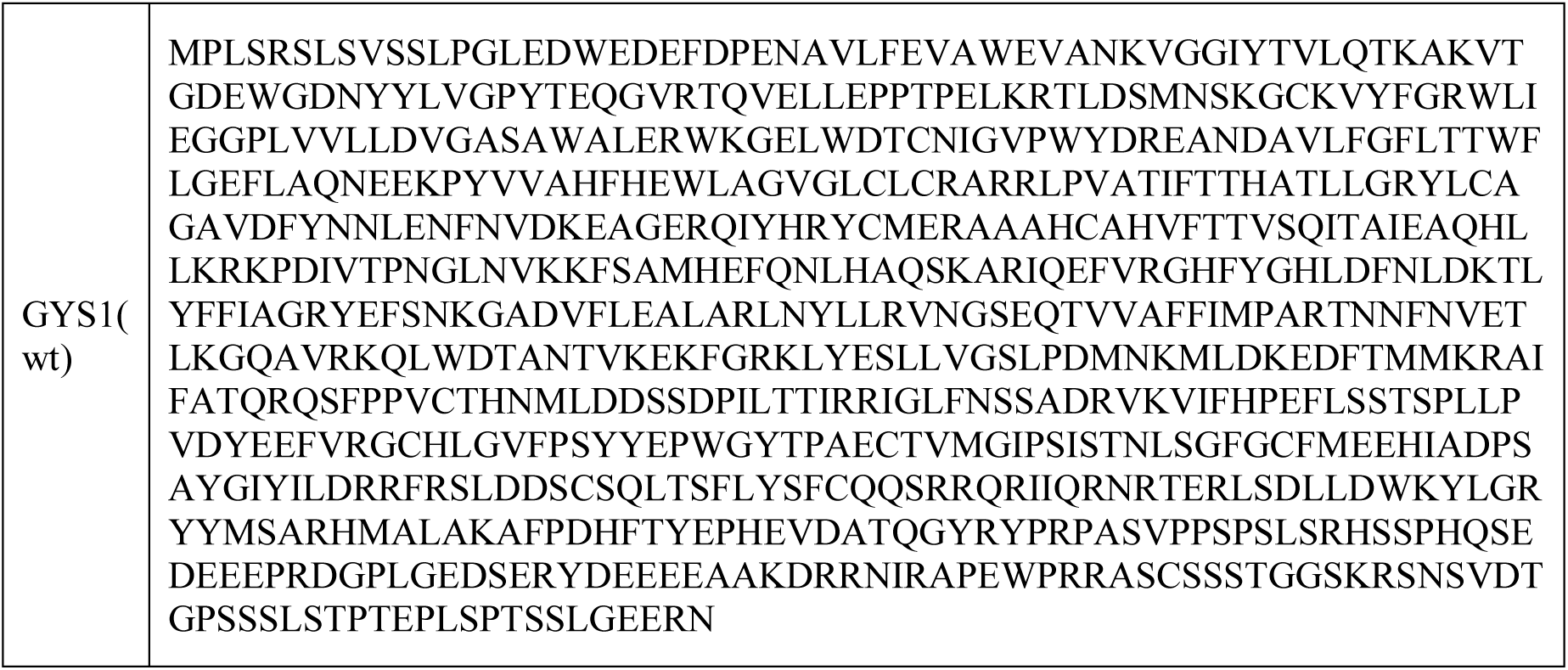
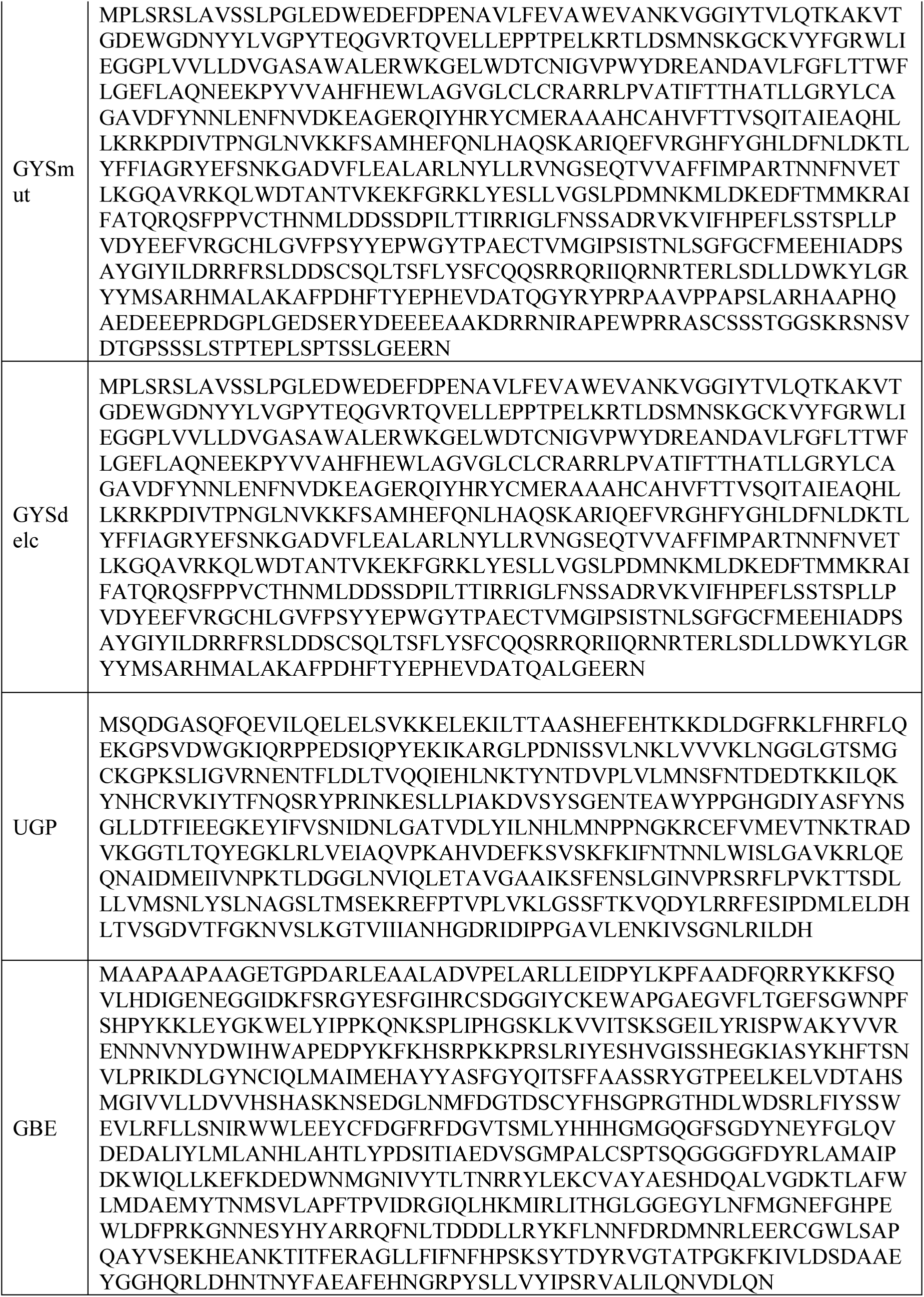
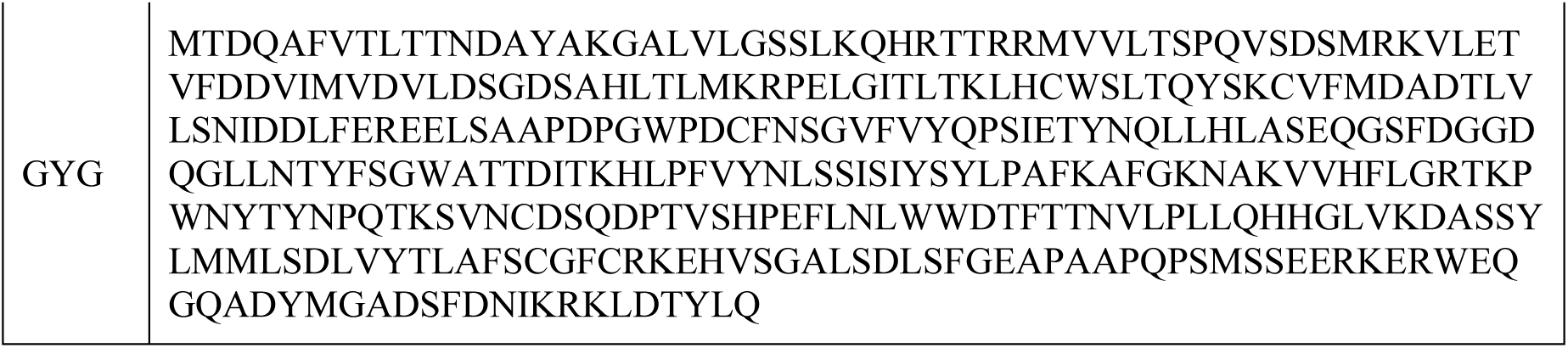
Protein sequences used in this study.

## References

1. Alonso-Gonzalez A, Tosco-Herrera E, Molina-Molina M, et al. Idiopathic pulmonary fibrosis and the role of genetics in the era of precision medicine. Front Med (Lausanne) 2023; 10: 1152211. 2023/05/14. DOI: 10.3389/fmed.2023.1152211.

2. Bari E, Ferrarotti I, Saracino L, et al. Mesenchymal Stromal Cell Secretome for Post-COVID-19 Pulmonary Fibrosis: A New Therapy to Treat the Long-Term Lung Sequelae? Cells 2021; 10 20210514. DOI: 10.3390/cells10051203.

3. Li Z, Niu S, Guo B, et al. Stem cell therapy for COVID-19, ARDS and pulmonary fibrosis. Cell Prolif 2020; 53: e12939. 20201024. DOI: 10.1111/cpr.12939.

4. Li DY, Li RF, Sun DX, et al. Mesenchymal stem cell therapy in pulmonary fibrosis: a meta-analysis of preclinical studies. Stem Cell Res Ther 2021; 12: 461. 20210818. DOI: 10.1186/s13287-021-02496-2.

5. Cheng SL, Lin CH and Yao CL. Mesenchymal Stem Cell Administration in Patients with Chronic Obstructive Pulmonary Disease: State of the Science. Stem Cells Int 2017; 2017: 8916570. 20170220. DOI: 10.1155/2017/8916570.

6. Rahman M, Wang ZY, Li JX, et al. Single-Cell RNA Sequencing Reveals the Interaction of Injected ADSCs with Lung-Originated Cells in Mouse Pulmonary Fibrosis. Stem Cells Int 2022; 2022: 9483166. 2022/04/23. DOI: 10.1155/2022/9483166.

7. Zhao X, Wu J, Yuan R, et al. Adipose-derived mesenchymal stem cell therapy for reverse bleomycin-induced experimental pulmonary fibrosis. Sci Rep 2023; 13: 13183. 20230814. DOI: 10.1038/s41598-023-40531-9.

8. Tian X, Jia Y, Guo Y, et al. Fibroblast growth factor 2 acts as an upstream regulator of inhibition of pulmonary fibroblast activation. FEBS open bio 2023; 13: 1895–1909. 2023/08/16. DOI: 10.1002/2211-5463.13691.

9. Moya A, Paquet J, Deschepper M, et al. Human Mesenchymal Stem Cell Failure to Adapt to Glucose Shortage and Rapidly Use Intracellular Energy Reserves Through Glycolysis Explains Poor Cell Survival After Implantation. Stem Cells 2018; 36: 363–376. 20180109. DOI: 10.1002/stem.2763.

10. Levy O, Kuai R, Siren EMJ, et al. Shattering barriers toward clinically meaningful MSC therapies. Sci Adv 2020; 6: eaba6884. 20200722. DOI: 10.1126/sciadv.aba6884.

11. Ladurner AG. Rheostat control of gene expression by metabolites. Mol Cell 2006; 24: 1–11. DOI: 10.1016/j.molcel.2006.09.002.

12. Lavrentieva A, Majore I, Kasper C, et al. Effects of hypoxic culture conditions on umbilical cord-derived human mesenchymal stem cells. Cell Commun Signal 2010; 8: 18. 20100716. DOI: 10.1186/1478-811x-8-18.

13. Das R, Jahr H, van Osch GJ, et al. The role of hypoxia in bone marrow-derived mesenchymal stem cells: considerations for regenerative medicine approaches. Tissue Eng Part B Rev 2010; 16: 159–168. DOI: 10.1089/ten.TEB.2009.0296.

14. Salazar-Noratto GE, Luo G, Denoeud C, et al. Understanding and leveraging cell metabolism to enhance mesenchymal stem cell transplantation survival in tissue engineering and regenerative medicine applications. Stem Cells 2020; 38: 22–33. 20191025. DOI: 10.1002/stem.3079.

15. Deschepper M, Manassero M, Oudina K, et al. Proangiogenic and prosurvival functions of glucose in human mesenchymal stem cells upon transplantation. Stem Cells 2013; 31: 526–535. DOI: 10.1002/stem.1299.

16. Roach PJ, Depaoli-Roach AA, Hurley TD, et al. Glycogen and its metabolism: some new developments and old themes. Biochem J 2012; 441: 763–787. DOI: 10.1042/bj20111416.

17. Jørgensen SB, Nielsen JN, Birk JB, et al. The alpha2-5’AMP-activated protein kinase is a site 2 glycogen synthase kinase in skeletal muscle and is responsive to glucose loading. Diabetes 2004; 53: 3074–3081. DOI: 10.2337/diabetes.53.12.3074.

18. Li Y, Wang T, Li X, et al. SOD2 promotes the immunosuppressive function of mesenchymal stem cells at the expense of adipocyte differentiation. Mol Ther 2024; 32: 1144–1157. 20240203. DOI: 10.1016/j.ymthe.2024.01.031.

19. Szwed A, Kim E and Jacinto E. Regulation and metabolic functions of mTORC1 and mTORC2. Physiol Rev 2021; 101: 1371–1426. 20210218. DOI: 10.1152/physrev.00026.2020.

20. Bozec D, Sattiraju A, Bouras A, et al. Akaluc bioluminescence offers superior sensitivity to track in vivo glioma expansion. Neurooncol Adv 2020; 2: vdaa134. 20201010. DOI: 10.1093/noajnl/vdaa134.

21. Huang K, Liu X, Qin H, et al. FGF18 encoding circular mRNA-LNP based on glycerolipid engineering of mesenchymal stem cells for efficient amelioration of osteoarthritis. Biomater Sci 2024; 12: 4427–4439. 20240820. DOI: 10.1039/d4bm00668b.

22. Contreras-Lopez RA, Elizondo-Vega R, Torres MJ, et al. PPARβ/δ-dependent MSC metabolism determines their immunoregulatory properties. Sci Rep 2020; 10: 11423. 20200710. DOI: 10.1038/s41598-020-68347-x.

23. Luo G, Wosinski P, Salazar-Noratto GE, et al. Glucose Metabolism: Optimizing Regenerative Functionalities of Mesenchymal Stromal Cells Postimplantation. Tissue Eng Part B Rev 2023; 29: 47–61. 20230125. DOI: 10.1089/ten.TEB.2022.0063.

24. Allen GM, Frankel NW, Reddy NR, et al. Synthetic cytokine circuits that drive T cells into immune-excluded tumors. Science 2022; 378: eaba1624. 20221216. DOI: 10.1126/science.aba1624.

25. Cheng S, Nethi SK, Rathi S, et al. Engineered Mesenchymal Stem Cells for Targeting Solid Tumors: Therapeutic Potential beyond Regenerative Therapy. J Pharmacol Exp Ther 2019; 370: 231–241. 20190607. DOI: 10.1124/jpet.119.259796.

26. Ye L, Park JJ, Peng L, et al. A genome-scale gain-of-function CRISPR screen in CD8 T cells identifies proline metabolism as a means to enhance CAR-T therapy. Cell Metab 2022; 34: 595–614.e514. 20220310. DOI: 10.1016/j.cmet.2022.02.009.

27. Tzouvelekis A, Koliakos G, Ntolios P, et al. Stem cell therapy for idiopathic pulmonary fibrosis: a protocol proposal. J Transl Med 2011; 9: 182. 20111021. DOI: 10.1186/1479-5876-9-182.

28. Crowley G, Kwon S, Caraher EJ, et al. Quantitative lung morphology: semi-automated measurement of mean linear intercept. BMC Pulm Med 2019; 19: 206. 20191109. DOI: 10.1186/s12890-019-0915-6.

